# Contribution of segment 3 to the acquisition of virulence in contemporary H9N2 avian influenza viruses

**DOI:** 10.1101/2020.05.27.119917

**Authors:** Anabel L. Clements, Joshua E. Sealy, Thomas P. Peacock, Jean-Remy Sadeyen, Saira Hussain, Samantha J. Lycett, Holly Shelton, Paul Digard, Munir Iqbal

## Abstract

H9N2 avian influenza viruses circulate in poultry throughout much of Asia, the Middle East and Africa. These viruses cause huge economic damage to poultry production systems and pose a zoonotic threat both in their own right as well as in the generation of novel zoonotic viruses, for example H7N9. In recent years it has been observed that H9N2 viruses have further adapted to poultry, becoming more highly transmissible and causing higher morbidity and mortality. Here, we investigate the molecular basis for this increased virulence, comparing a virus from the 1990s and a contemporary field strain. The modern virus replicated to higher titres in various systems and this difference mapped to a single amino acid polymorphism at position 26 of the endonuclease domain shared by the PA and PA-X proteins. This change was responsible for the virulent phenotype and extended tissue tropism seen in chickens. Although the PA K26E change correlated with increased host cell shutoff activity of the PA-X protein *in vitro*, it could not be overridden by frameshift site mutations that block PA-X expression and therefore increased PA-X activity could not explain the differences in replication phenotype. Instead, this indicates these differences are due to subtle effects on PA function. This work gives insight into the ongoing evolution and poultry adaptation of H9N2 and other avian influenza viruses and helps us understand the soaring morbidity and mortality rates in the field, as well as rapidly expanding geographical range seen in these viruses.

**Author Summary:** Avian influenza viruses, such as H9N2, cause huge economic damage to poultry production worldwide and are additionally considered potential pandemic threats. Understanding how these viruses evolve in their natural hosts is key to effective control strategies. In the Middle East and South Asia an older H9N2 virus strain has been replaced by a new reassortant strain with greater fitness. Here we take representative viruses and investigate the genetic basis for this ‘fitness’. A single mutation in the virus was responsible for greater fitness, enabling high growth of the contemporary H9N2 virus in cells, as well as in chickens. The genetic mutation that modulates this change is within the viral PA protein, a part of the virus polymerase gene that contributes in viral replication as well as contribute in the virus accessory functions – however we find that the fitness effect is specifically due to changes in the protein polymerase activity.

## Introduction

Influenza A viruses possess a segmented, negative-sense RNA genome which is transcribed and replicated by a tripartite RNA-dependent RNA polymerase (RdRp) composed of the subunits PB2, PB1 and PA. Due to the segmented nature of its genome, influenza viruses can readily swap genes when two virus strains co-infect a single cell, in a process known as reassortment. Reassortment can result in generation of viruses with increased [1] or reduced viral fitness [2].

H9N2 avian influenza viruses (AIVs) are low pathogenicity avian influenza (LPAI) viruses that are enzootic in poultry in many countries across Asia, Africa and the Middle East [3-6]. In afflicted countries they cause a constant burden on poultry production systems through mortality often associated with co-infection, or morbidity that leads to reduced egg production and bird growth rates [7-9]. They also pose a zoonotic risk as evidenced by over 60 confirmed cases of human infection, with over half of those occurring since 2015 [6].

Due to their extensive geographical range H9N2 AIVs often co-circulate with other AIV subtypes resulting in frequent reassortment events [10]. Several viruses have emerged in recent years which contain the internal gene cassette derived from H9N2 AIVs including a novel avian-origin H7N9 virus which has caused human infections in China. H7N9 possesses the polymerase genes from an enzootic co-circulating H9N2 strain [11, 12]. Novel genotypes of H9N2 AIV have also emerged in poultry due to co-circulation and reassortment with local highly pathogenic avian influenza virus strains; we have previously described G1-lineage H9N2 viruses in Pakistan that possess the NS gene segments from H7N3 or H5N1 strains and the polymerase genes from other Indian/Middle East lineage H9N2 viruses [13]. These reassortants have replaced previously circulating genotypes of the G1-lineage H9N2 AIVs, and are now the predominant genotype across the Indian subcontinent and Middle East, and display enhanced morbidity and mortality in the field [14-16].

The molecular basis for the increased pathogenicity of contemporary reassortant H9N2 AIVs has yet to be established. Thus, we set out to understand which genes are responsible for the enhanced virulence of these H9N2 viruses in poultry. We created a panel of reverse genetics reassortants between a pair of G1-lineage viruses A/guinea fowl/Hong Kong/WF10/1999 (WF10), a virus representing G1-lineage viruses circulating in the late 1990s, and A/chicken/Pakistan/UDL-01/2008 (UDL-01), representative of novel G1-lineage reassortant H9N2 viruses. UDL-01 contains the HA, NA, NP and M genes related to previously circulating enzootic G1-lineage H9N2 viruses in the region, the polymerase gene cassette from different G1-lineage H9N2 viruses, more predominant in the Middle East, and the NS gene segments from an HPAIV H7N3 [13].

In this study we find that the ancestral virus WF10, showed an attenuated replication phenotype *in vitro* when compared to the contemporary H9N2 virus, UDL-01. This phenotypic difference mapped to a single amino-acid residue in the PA endonuclease domain (position 26) within segment 3. This single residue also determined the replicative fitness and virulence of the virus *in vivo*, and was further shown to modulate the activity of PA in a PA-X independent manner.

## Results

### Differences in plaque phenotype between two H9N2 AIV strains maps to the N-terminal half of segment 3

We generated a panel of reciprocal reassortant viruses between the full reverse genetics systems of WF10, a virus representing G1-lineage H9N2 AIVs that circulated in the late 1990s, and UDL-01, representative of a novel reassortant G1-lineage H9N2 with genes from several previously enzootic G1-lineage H9N2 viruses and HPAIV H7N3 viruses. Wild-type (WT) WF10 virus generated small hazy plaques in MDCK cells whereas WT UDL-01 generated significantly larger, clearer plaques (Figure 1A, B). We tested the plaque phenotype of all reassortants and identified segment 3 as capable of reciprocating plaque phenotype between WF10 and UDL-01 (Figure 1A, B; data for non-segment 3 reassortants not shown). UDL-01 virus containing segment 3 of WF10 presented significantly smaller plaques, while reassortant WF10 presented significantly larger plaques, relative to WT viruses (Figure 1A, B).

**Figure 1.**
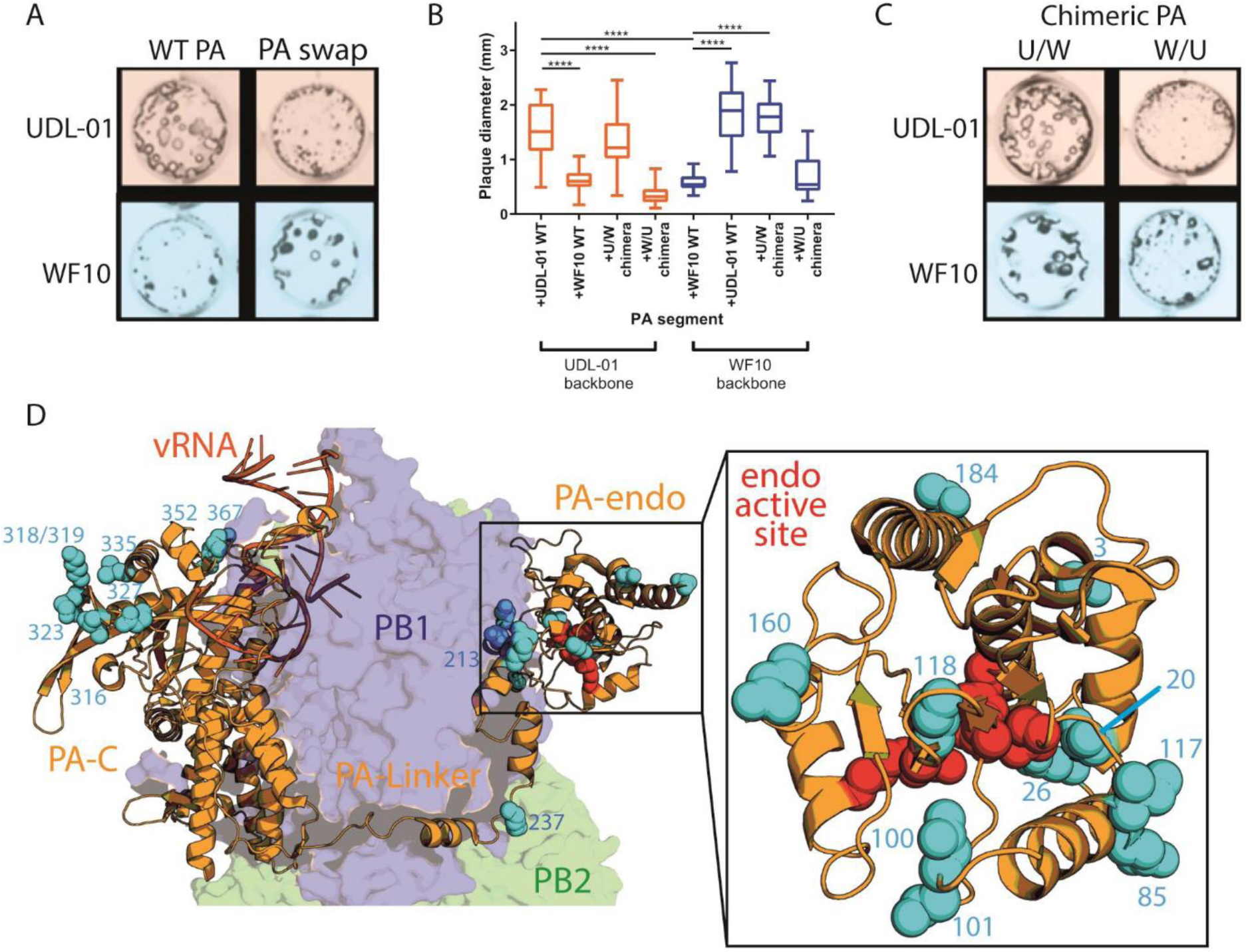
The small plaque phenotype of WF10 maps to the N-terminus of segment 3. The indicated viruses were titrated in MDCK cells via plaque assay under a 0.6% agarose overlay and 72 h.p.i., fixed and stained for NP using immunofluorescence. (A) and (C) show representative images of plaque sizes of UDL-01 and WF10 RG viruses. (B) shows diameter of 20 plaques/virus measured using Image J analysis software. The graphs represent the average plaque diameter +/- SD. (B) Kruskal-Wallis with Dunn’s multiple comparison test was used to determine the statistical differences between the plaque sizes. **** P value < 0.0001 (D) Structure of the trimeric polymerase with vRNA (dark orange) with PB2 (green), PB1 (blue) and PA (light orange).N-terminus half PA differences between WF10 and UDL-01 shown in cyan, zoomed in PA endonuclease domain included in right panel, PA endonuclease active set residues H41, E80, D108, E119 and K134 shown in red (PDB ID: 4WSB)[27].

To identify the region of segment 3 responsible for this alteration, two chimeric segments 3s were generated by Gibson assembly: one chimera encoded the N-terminus (amino acids 1-367) from PA of UDL-01 and the C-terminus (amino acids 368-716) of WF10 (U/W), while the other was *vice versa* (W/U). These chimeric segments were rescued by reverse genetics in the background of UDL-01 and WF10 viruses. Determination of the plaque phenotypes showed that, regardless of the rest of the virus genes, viruses containing a UDL-01 PA N-terminal coding region had significantly larger plaques that those without (Figure 1B, C). Furthermore, UDL-01 virus, which typically presents a large plaque phenotype, presented significantly smaller plaques when given a WF10 N-terminus (Figure 1B, C). These results show that the small plaque phenotype could be mapped specifically to the N-terminal half of WF10 PA.

### Amino acid residue 26 in PA modulates plaque phenotype

PA is composed of two major domains, an N-terminal endonuclease (endo) domain and a C-terminal domain (PA-C) connected by a linker region [17]. The PA endo domain is a flexible appendage that hangs away from the catalytic core of the RdRp and is involved in cleaving host capped RNAs to be fed into the RdRp active site be used as primers for viral transcription. The PA-C domain is packed close to PB1 and makes up part of the catalytic core of the viral RdRp [18].

The first 191 amino acids of PA, incorporating the endo domain, are also shared with the accessory protein PA-X. PA-X is expressed due to a ribosomal frame shift site in PA, during segment 3 translation a small proportion of ribosomes, when encountering a rare tRNA codon slip into the +2 open reading frame (ORF) of segment 3 and express a fusion protein comprised of the PA endo domain and an X-ORF from the +2 reading frame [19, 20]. PA-X dampens the innate immune response though its host cell shutoff activity, mediated through degradation of cellular mRNAs and disruption of mRNA processing machinery [19, 21]. The evidence for PA-X playing a role as a virulence factor in avian influenza viruses is unclear, with several studies showing either attenuation or promotion of virulence *in vivo* [22-26].

To identify amino acid substitutions responsible for modulating virus plaque phenotype, we compared amino acids in the N-terminal half of PA between UDL-01 and WF10, as well as between the X-ORFs of PA-X. We identified a total of twenty amino acid differences between PA and 2 unique to the X-ORF (Table 1). Mapping the residues onto the crystal structure of influenza PA within the context of the polymerase trimer bound to vRNA showed that they lay in the PA-endo domain, the linker region and at the N-terminus of the PA-C domain [27] (Figure 1D, Table 1). This enabled us to speculate if any of the substitutions had a direct effect due to their proximity to known functional regions. Substitution I118T is specifically located within the endonuclease active site, while A20T, E26K and I100V/D101E lie proximal to the active site and could potentially interfere with endonuclease activity. Therefore, these mutations, alongside the X-ORF polymorphisms and several other mutants at residues shown previously to modulate polymerase activity [28], were selected for further testing.

**Table 1:** Amino acid differences between the N-terminus of progenitor (WF10) and reassortant (UDL-01) H9N2 segment 3 products.

| Amino acid position | WF10 | UDL-01 | Domain |
| --- | --- | --- | --- |
| 3 | D | N | Endo |
| 20 | T | A | Endo |
| 26 | K | E | Endo |
| 85 | A | T | Endo |
| 86 | M | L | Endo |
| 100 | V | I | Endo |
| 101 | D | E | Endo |
| 118 | T | I | Endo |
| 160 | D | E | Endo |
| 184 | S | N | Endo |
| 213 | R | K | Linker |
| 237 | K | E | Linker |
| 316 | D | G | PA-C |
| 318 | R | K | PA-C |
| 319 | E | D | PA-C |
| 323 | I | V | PA-C |
| 327 | E | K | PA-C |
| 335 | I | L | PA-C |
| 352 | D | E | PA-C |
| 367 | M | K | PA-C |
| X-221 | L | R | X-ORF (PA-X) |
| X-250 | R | Q | X-ORF (PA-X) |

A panel of viruses was made carrying reciprocal single amino acid substitutions at the sites identified and viruses were rescued using reverse genetics. Within the UDL-01 panel of PA mutant viruses, two mutants gave significantly smaller plaque sizes than UDL-01 WT: the E26K mutant produced comparably-sized plaques to that of the WF10 WT virus (Figure 2A,B,C), while the double mutant I100V/E101D also had a small plaque phenotype, though less markedly so than E26K (Figure 2C). Within the WF10 panel of viruses, the most striking visual difference was caused by the introduction of K26E, which facilitated significant larger plaque diameters (Figure 2B, C). D316G also gave a heterogenous but significantly larger average plaque size than WT WF10. Mutations at position 26 were the only viruses to give reciprocal plaque size phenotypes in both viral backgrounds, strongly suggesting that this position is key to the phenotype.

**Figure 2.**
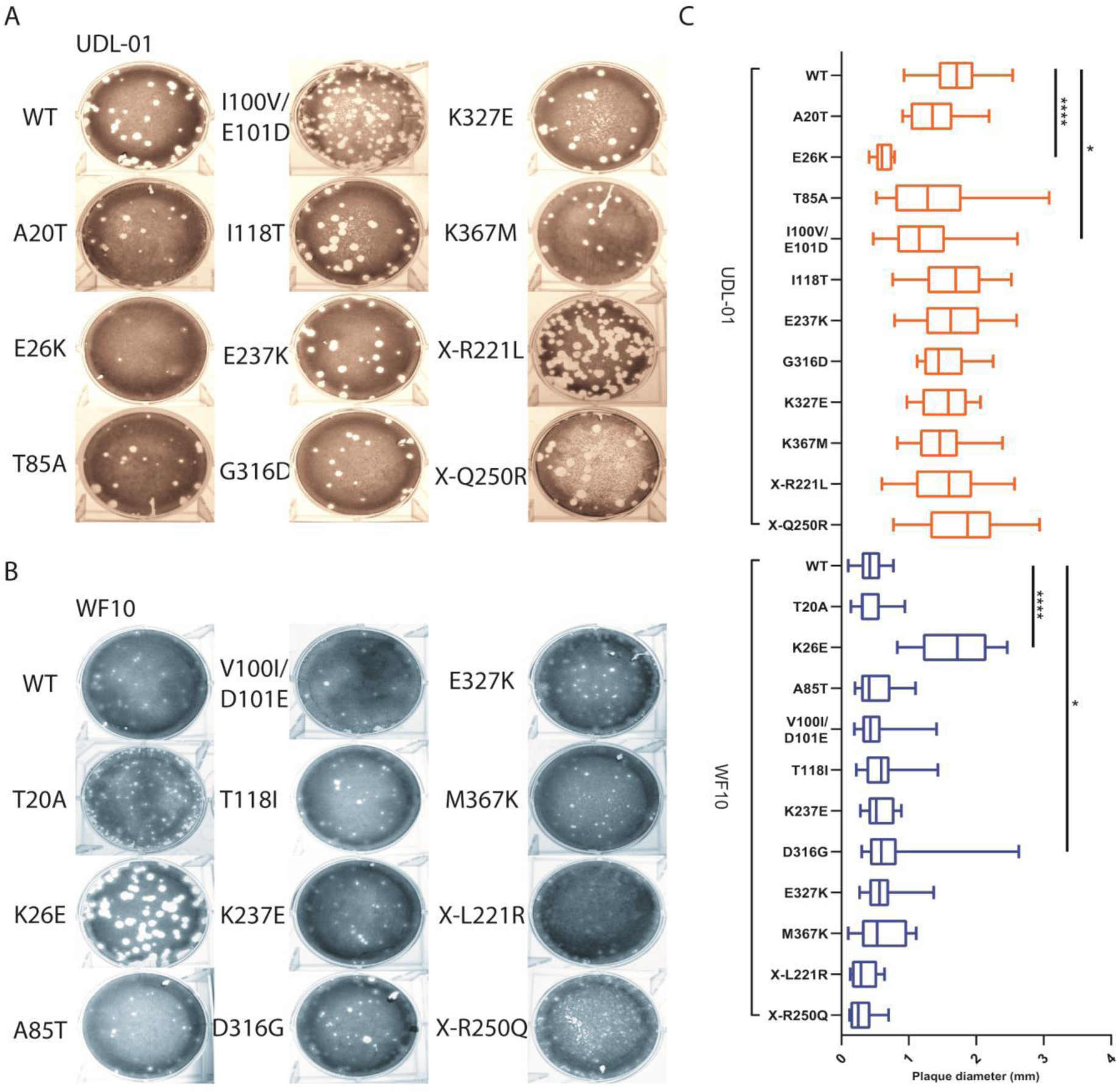
The small plaque phenotype of WF10 maps to PA position 26. H9N2 AIVs with either a UDL-01 or WF10 backbone were rescued via reverse genetics. The plaque phenotypes of the rescues were assessed via plaque assay on MDCK cells under a 0.6% agarose overlay. After 72 hours cells were fixed and stained with 0.1% crystal violet solution and plaques imaged. (A) Visual representation of UDL-01 virus panel containing PA mutations to make them WF10-like. (B) Visual representation of WF10 virus panel containing PA mutations to make them UDL-01 like. (C) 20 plaque diameters per virus performed on the same day were measured using ImageJ analysis software and the average plaque diameter calculated. Graph represents the average +/- SD. P values= ****: <0.0001; **: <0.0039 (Kruskal-Wallis with Dunn’s multiple comparisons).

### Residue 26 in PA modulates virus replication kinetics

For influenza viruses, small plaque phenotypes are often used as a marker of poor virus replication, thus we further investigated this phenotype by performing multiple cycle replication kinetics experiments with the position 26 mutants.

In MDCK cells infected at a low MOI, by later time points (36 hours and after) UDL-01 WT clearly grew to higher titres than WF10 WT, consistent with the plaque assay phenotypes (Figure 3A). UDL-01 E26K showed slightly attenuated growth compared to UDL-01 WT while WF10 K26E showed slightly enhanced titres compared to WT WF10, which were significantly different at the 48 and 72 hour time points (Figure 3A). Thus, PA residue 26 had significant reciprocal effects on the replication of UDL-01 and WF10 viruses in MDCK cells, recapitulating the differences seen for the MDCK plaque size phenotype.

**Figure 3.**
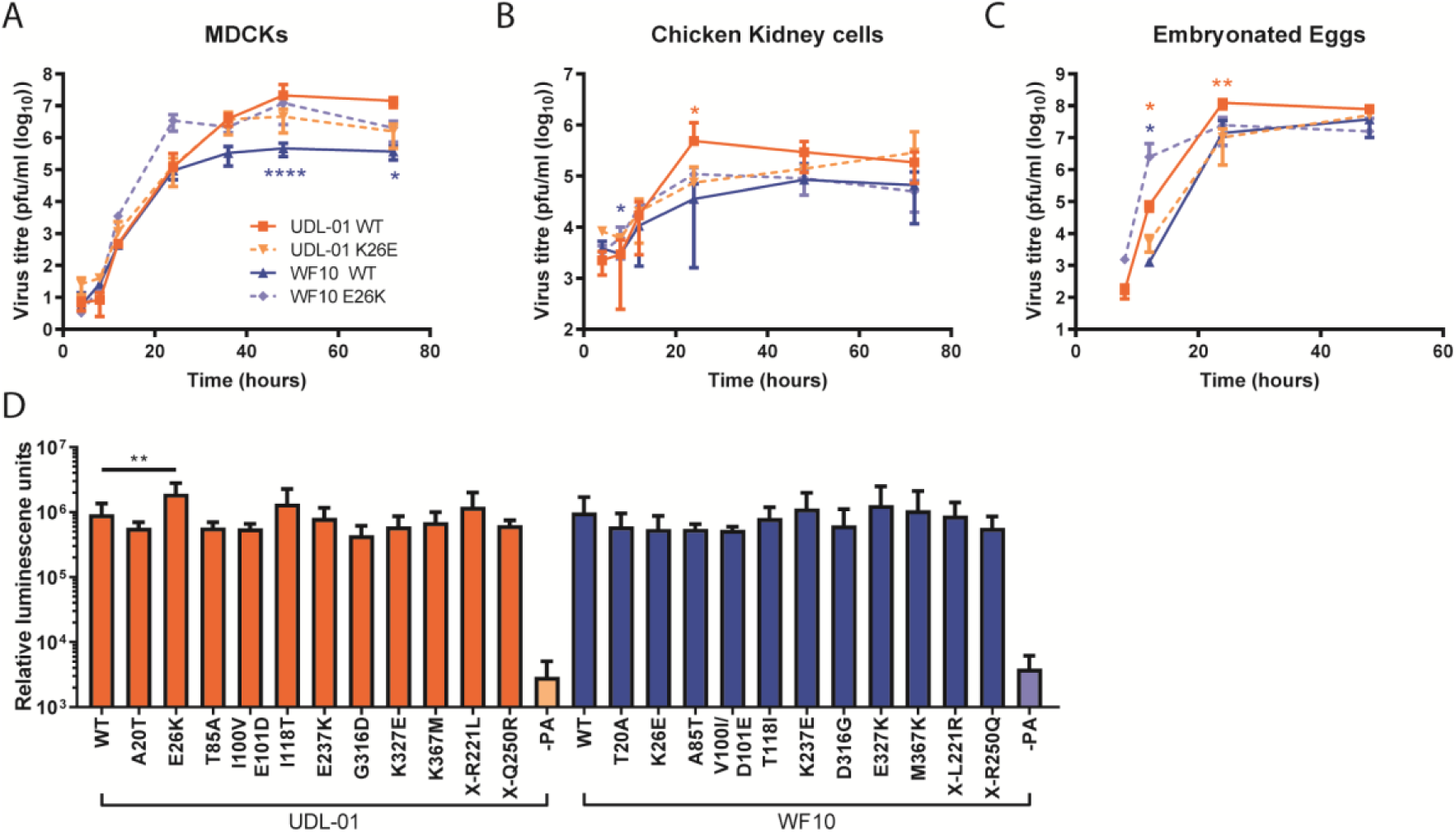
Variation at position 26 leads to differences in replication but not polymerase activity. (A) MDCK cells and (B) CK cells were infected with the specified virus (UDL-01 WT, UDL-01 E26K, WF10 WT or WF10 K26E) at a low MOI (0.01). (C) 10-day-old fertilised hens’ eggs infected with 100pfu of virus. Samples were taken at the indicated time points for titration via plaque assay. No virus was detected prior to 8 h.p.i. Data represents the average +/- SD of 3 independent experiments (cells) or 5 eggs per timepoint. Significant differences (unpaired T-tests (A; UDL-01 36 and 72 h.p.i. WF10 24, 36, 48 and 72 h.p.i., B: UDL-01 8, 12, 24 and 48 h.p.i. WF10 8, 48 and h.p.i. C: UDL-01 12 h.p.i.) or Mann-Whitney Test: (A; UDL-01 4, 8, 12, 24 and 48 h.p.i. WF10 4, 8 and 12 h.p.i. B; UDL-01 4 and 72 h.p.i. and WF10 4, 12 and 72 h.p.i. C: UDL-01 8, 24 and 48 h.p.i. WF10 all data points) between WT and corresponding mutant at each time point depending on distribution of data are represented via with asterisks in orange (UDL-01 pair) or blue (WF10 pair). P values: *= <0.035; ** = <0.008; **** = <0.0001. (D) Polymerase activity of the different mutants was assessed using an influenza minireplicon assay. DF-1 cells were transfected with the components of the polymerase complex (PB1, PB2, PA and NP) plus a vRNA mimic encoding luciferase under the control of an avian RNA polymerase I promotor. 48 hours post transfection cells were lysed and luciferase levels measured. Data represents the mean +/- SD of 3 independent experiments.

To test if the effect of PA residue 26 amino acid substitutions held true in more biologically relevant avian systems, viral replication kinetics were assessed in primary chicken kidney (CK) cells and embryonated chicken eggs (Figure 3B, C). In CK cells there were consistent differences between the replication kinetics of the viruses similar to that seen in MDCKs; viruses with PA 26E (UDL-01 WT and WF10 K26E) reached peak titres at 24 hours post-infection, while 26K-containing viruses replicated at a slower rate, achieving maximum titres at 48 and 72 hour time points (Figure 3B). UDL-01 E26K trended towards lower titres than UDL-01 WT, which was significant at 24 hours post-infection. Likewise, WF10 K26E generally showed enhanced titres compared to WF10 WT, which was significant at 8 hours post-infection.

In embryonated eggs, as in MDCK cells and CK cells, UDL-01 E26K showed attenuated growth compared to UDL-01 WT, significantly so at 12 and 24 hours post-infection, while WF10 K26E showed enhanced growth compared to WF10 WT, significantly at 12 hours post-infection (Figure 3C). Considering these data together, we can conclude that amino acid substitutions at position 26 of PA within WF10 and UDL-01 H9N2 AIVs significantly altered the replication of the viruses in both mammalian cell lines, and avian systems indicating this attenuation is not host-dependent.

### Impact of PA amino acid substitutions on polymerase activity

As an integral part of the trimeric polymerase, influenza PA mutations have previously been shown to impact polymerase activity due to its position within the heterotrimeric polymerase complex (*e*.*g*. [29]). To investigate the role of K26E, as well as the other mutations tested here on polymerase activity, minireplicon assays were performed in chicken DF-1 cells. Cells were transfected with expression plasmids for either the UDL-01 or WF10 polymerase components plus NP and a vRNA-reporter encoding luciferase under the control of the avian RNA polymerase I promotor. No significant differences were seen between the activities of polymerase complexes containing UDL-01 or WF10 WT polymerases (Figure 3D). However, in contrast to the virus replication assays, UDL-01 E26K showed a small (∼ 2-fold) but significant increase in polymerase activity compared to UDL-01 WT. There were no further significant differences seen with any of the UDL-01 or WF10 mutants when compared to the activity of the relevant WT control, including the reciprocal K26E change in WF10. Polymerase activity was also measured in mammalian 293T cells but no significant differences were seen (data not shown). These data suggest that the differences in plaque phenotype observed between WF10 (progenitor) and UDL-01 (reassortant) H9N2 AIVs was not unambiguously related to the segment 3s ability to support polymerase activity alone..

### PA-E26K attenuates virus replication and pathogenicity *in vivo*

The WF10-like K26E mutation appears to lead to an attenuated replication phenotype for UDL-01 *in vitro, ex vivo* and *in ovo*, therefore we decided to assess the ability of UDL-01 E26K within the natural chicken host. Two groups of chickens were inoculated with either UDL-01 WT or UDL-01 E26K virus and viral shedding, transmissibility, tissue tropism and clinical signs were assessed. In both infected groups all directly inoculated birds shed virus robustly into the buccal cavity, peaking early in infection (day 1 or 2) and then declining over the subsequent days (Figure 4A). However, the UDL-01 E26K virus was shed in significantly lower amounts and was cleared sooner; no swabs were found positive for infectious virus by day 5 in the E26K group compared to day 7 for the WT group (Figure 4A). When the area under the shedding curves (AUC) were calculated to assess the total virus shed throughout the study period, birds directly infected with UDL-01 WT virus showed almost a ten-fold increased AUC compared to birds infected with the mutant UDL-01 E26K virus (215,517 versus 23,886). Therefore, the mutant UDL-01 E26K virus showed a reduced total shedding throughout the study by birds directly infected with virus.

**Figure 4.**
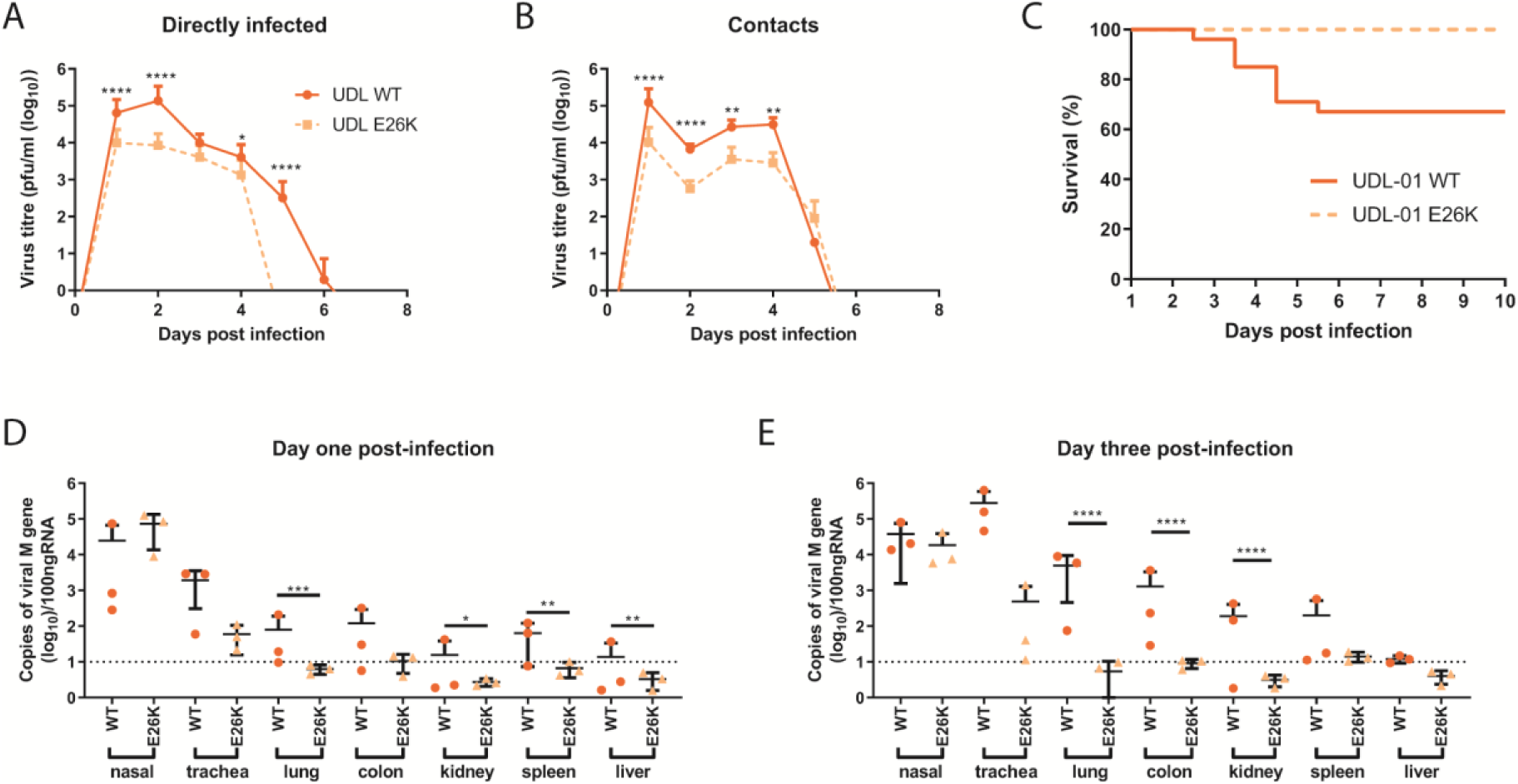
In the background of UDL-01, K26E leads to reduced shedding, mortality and tissue tropism. Groups of 20 chickens were infected with 10^4^pfu of either UDL-01 WT or mutant UDL-01 E26K viruses. One day post-infection 8 naïve contact birds were cohoused with each group. Birds were swabbed in the buccal and cloacal cavities throughout the study duration. (A, B) average buccal shedding profile of directly infected or contact birds. (C) Survival curve of birds exposed to each virus, graph includes both birds that died spontaneously or which reached a humane end point (D, E) qRT-PCR for detection of M gene of viral RNA from chicken tissues of birds culled on day 1 and 3 post-infection. Mann-Whitney Test was conducted for A and B. For D and E, unpaired T-test (Day 1-Trachea, Kidney, Spleen and Liver, Day 3-Nasal, Trachea, Colon, Kidney, Spleen) or Mann Whitney Tests (Day 1-Lung and Colon, Day 3-Lung and Liver) between UDL-01 WT and UDL-01 E26K infected birds was conducted. P values = ****:<0.001, ***: <0.0005, **: <0.0082, *:<0.033. Error bars represent +/- SD of all birds swabbed (at least 4 birds/group). For survival curves (E) P value = <0.0001 (Log rank (Mantel-Cox) test).

Contact birds were introduced into each group 1 day post-inoculation. All contact birds in both groups tested positive for infectious virus from the buccal cavity by 1 day post-exposure indicating robust contact transmission for both viruses (Figure 4B). A significant reduction in buccal shedding was seen in contact birds exposed to UDL-01 E26K compared to UDL-01 WT from day 1 through to 4 post-exposure; however, the delayed clearance of the mutant virus was not seen in the contact bird group, with both groups of birds clearing virus by day 6 post-exposure and similar levels of virus being shed on day five post-exposure (Figure 4B). When the AUC was calculated to assess the total virus shed throughout the study period, contact birds infected with UDL-01 WT virus again showed around a ten-fold greater AUC compared to birds infected with the mutant UDL-01 E26K virus (187,915 versus 17,367).

Cloacal swabs from directly infected and contact birds were also analysed, shedding was sporadic with not all birds yielding detectable infectious virus (data not shown). In total, six UDL-01 WT and five E26K directly infected birds shed detectable virus along with a single contact bird from each group. More birds shed virus on consecutive days in the WT group than the E26K group (4 birds versus 1). This sporadic and low level virus shedding is commonly seen for some AIV subtypes including the UDL-01 virus [30-33].

Clinical signs throughout the study were generally mild and the majority of birds showed diarrhoea with depression as expected from previous reports of H9N2 infection, including UDL-01 H9N2 [30, 33]. However, between days three to six post-inoculation 33% of birds (30% directly infected and 37.5% of contact birds) within the UDL-01 WT infected group died (either spontaneously or due to reaching humane end points and being culled) despite UDL-01 being classified as a LPAIV (Figure 4C); analysis of these survival curves showed a statistically significant difference (p <0.0001). UDL-01 has previously shown to cause high levels or morbidity as well as occasionally low levels of mortality in experimentally infected animals of certain chicken lines [30, 33]. Birds infected with mutant UDL-01 E26K showed no mortality, indicating this single mutation clearly attenuates the virus for pathogenicity and mortality.

### Tropism of virus in infected chickens

To determine whether the UDL-01 E26K mutation lead to any alteration in tropism, tissues were taken from directly infected birds on days 1 and 3 post-inoculation. RNA extracted from tissue samples was used for qRT-PCR reactions to detect the viral M gene vRNA as a marker for presence of virus within tissues; which we have previously shown correlates well with tissue infectious virus titres [30]. At both time points, M gene copy number was highest in the nasal and tracheal tissues but was also readily detectable within the lung, colon, kidney, and spleen and intermittently detected in the liver (at least within UDL-01 WT infected birds; Figure 4D, E). Overall, RNA copies were variable between days, and between different birds, but levels were highest on day 3 post-inoculation, particularly within the visceral organs. Within these animals, UDL-01 WT virus was consistently present at higher levels in a number of tissues compared to birds infected with the mutant UDL-01 E26K virus. On day 1 post-inoculation there was significantly higher levels of RNA within the lung, kidney, spleen and liver of the UDL-01 WT, compared to UDL-01 E26K infected birds (Figure 4D). On day 3, UDL-01 E26K mutant virus was mostly undetectable in the visceral organs, but detectable within the nasal tissue where levels remained high (10^4^ to 10^5^ copies of viral M gene). The lung, colon and kidneys also showed significantly higher levels of UDL-01 WT RNA compared to the mutant UDL-01 E26K (Figure 4E). This suggested that although the UDL-01 E26K virus was able replicate efficiently in the upper respiratory tract, it was less able to disseminate through the bird and was more rapidly cleared. The expanded visceral tropism of the UDL-01 WT virus likely explains its highly mortality observed in this experimental study.

### Polymorphisms at position 26 affect the host shutoff activity of the accessory protein PA-X

As we observed little or no difference in polymerase activity with the reciprocal mutants at position 26 we hypothesized that the difference in replication could be due to WF10 having poor PA-X activity. To test this, a previously described β-galactosidase (β-gal) reporter assay was used to test the ability of the PA-X proteins from these viruses to cause host cell shutoff [19, 34]. Briefly, cells were co-transfected with expression plasmids containing the different segment 3s along with a β-gal reporter plasmid, followed by enzymatic readout of β-gal activity to give a measure of host gene expression in the transfected cells and thus the ability of the different segment 3 plasmids to cause host shutoff. Previous work has suggested that the majority of influenza host cell shutoff comes from expression of PA-X rather than PA [19, 34, 35].

Mammalian 293T cells were transfected with plasmids with or without mutations in the shared PA/PA-X endo domain or PA-X X-ORF and β-gal activity was measured. All data were normalized to a control where segment 3 was substituted for an empty vector. UDL-01 WT segment 3 significantly reduced levels of β-gal compared to the empty vector control indicating robust shutoff activity (Figure 5A), as shown previously [34]. In contrast WF10 WT segment 3 displayed no detectable shutoff, giving equivalent β-gal signal to the empty vector control. When the reciprocal mutants were tested, only mutations at position 26 had a reciprocal effect, significantly removing shutoff activity in a UDL-01 background and causing shutoff activity in the WF10 background. In the UDL-01 background I118T additionally showed significantly reduced shutoff activity but a reciprocal effect was not seen in WF10. In the WF10 background the X-ORF mutation X-L221R also significantly increased host shutoff activity. All other mutations showed the same phenotypes as their respective WT segment 3s.

**Figure 5.**
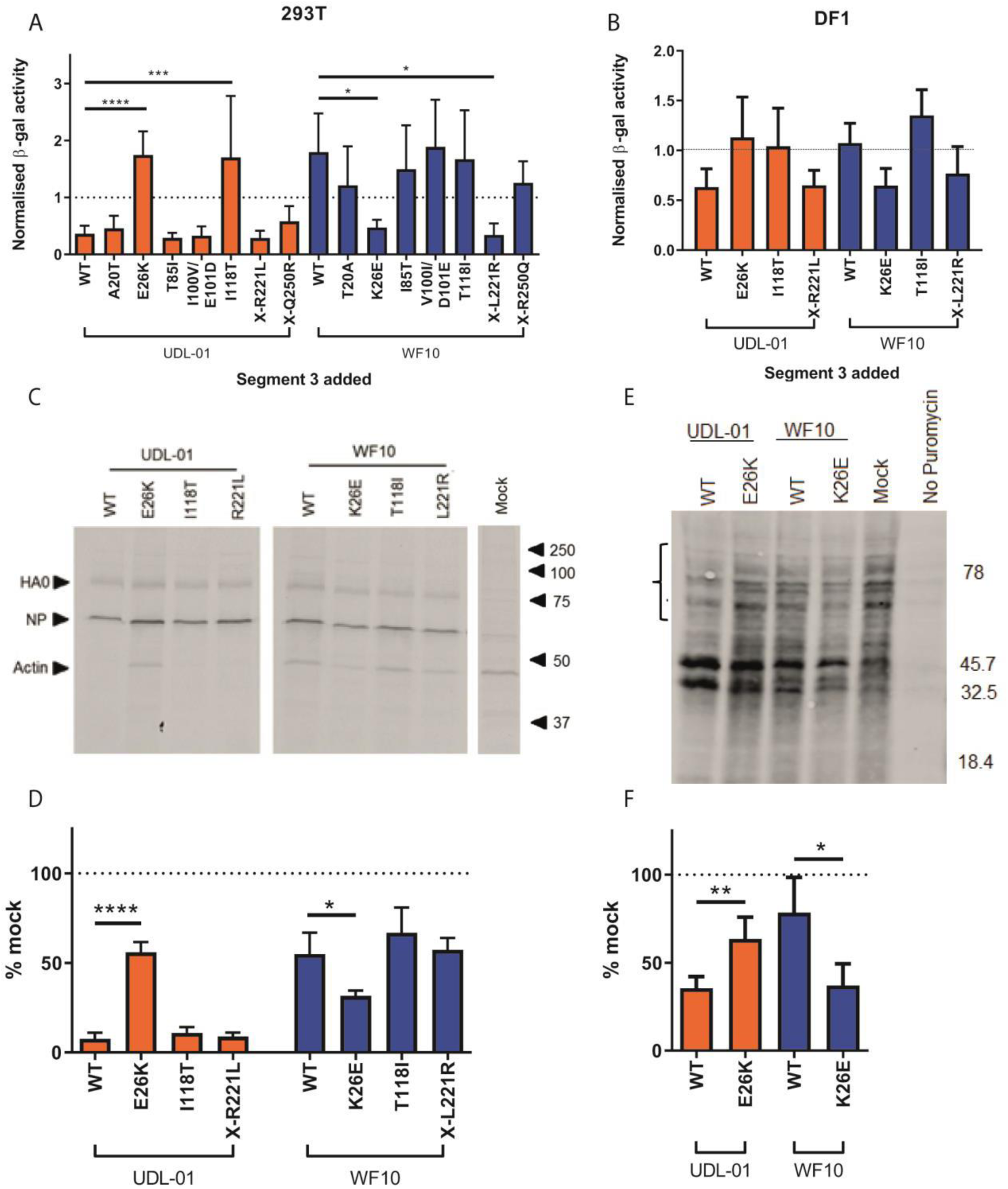
Lysine at position 26 correlates with a lack of host shutoff activity in PA-X. WT or mutant segment 3 plasmids were co-transfected into (A) 293T cells or (B) DF-1 cells with a β-gal reporter plasmid. 48 hours post-transfection cells were lysed and levels of β-gal assessed by colorimetric enzyme assay. Results were normalised to a sample where the PA plasmid was replaced with the empty vector control. Graph represents the average of 3 independent experiments +/- SD. P values: ****= <0.0001; ***= 0.0003; * = <0.003 (A, B) one-way ANOVA with multiple comparisons (all 293T and WF10 panel in DF-1) or Kruskal Wallis with multiple comparisons (UDL-01 panel in DF-1). (C, D) CEF were infected with a high MOI (3) of virus. 7 hours post-infection cells were pulsed with 35-S methionine for 1 hour then lysed and proteins separated by SDS-PAGE. Radiolabelled proteins were detected by autoradiography. (C) Representative SDS-PAGE gel with specific proteins and the positions of molecular mass (kDa) markers indicated. (D) Levels of radiolabelled actin were quantified by densitometry using ImageJ analysis software. Graph represents the average of 3 independent experiments +/- SD P values = *: 0.02, ****: <0.0001 (one-way ANOVA with multiple comparisons). (E, F) MDCK cells were infected with a high MOI (10) of each virus. 7.5 hours post-infection cells were pulsed with puromycin for 30 minutes. Cells were lysed, run on SDS-PAGE gels and western blotted for puromycin. (E) Representative western blot gel probed for puromycin. (F) The bracket in (E) covering the areas above 45.7kDa indicates the region quantified using ImageJ analysis software to measure the area under the curve following densitometry of this region. Data were converted to a percentage of the value seen in mock infected cells. Graph represents average +/- SD of 3 independent experiments. P values = *: 0.0124, **: 0.007 (unpaired T-test).

The host shutoff assay was also performed in avian DF-1 cells for the mutants at positions 26, 118 and X-221 (Figure 5B). Although not significant, the change at position 26 trended towards switching the two segment 3s phenotypes – removing shutoff activity from UDL-01 and introducing the activity into WF10 segment 3. Again UDL-01 I118T removed shutoff activity and WF10 X-L221R partially introduced shutoff activity, but as in 293Ts, these effects were not seen reciprocally.

To assess shutoff activity further, in the context of viral infection rather than transfection and overexpression, we performed both radioactive and non-radioactive metabolic labelling experiments. To test the shutoff activity in avian cells, primary chicken embryonic fibroblast (CEF) cells were infected with a high MOI of virus containing mutations in segment 3 and were subsequently pulsed with ^35^S methionine then lysed and run on SDS-PAGE. Autoradiography was performed and densitometry used to measure the abundance of the highly abundant host protein, actin. In accordance with the reporter assays, UDL-01 WT virus showed efficient host shutoff with <10% of the levels of actin expressed in the mock infected cells, whereas WF10 WT virus resulted in poor shutoff activity (>50% actin expressed versus mock; Figure 5C, D). Reciprocal mutants at position 26 were the only mutants tested that showed any significant effect; UDL-01 E26K showed significantly poorer host shutoff while WF10 K26E showed significantly more robust shutoff. Finally, a similar experiment was performed in MDCK cells using the non-radioactive method of puromycin pulsing and looking for levels of puromycinylated proteins in cell lysates [36]. MDCKs were either infected with WT viruses or the position 26 mutants. Levels of puromycinylated products were then detected by western blot and quantified in the ∼ 50 – 80 kDa range, where no novel products likely corresponding to viral polypeptides were visible. As with the previous assays, UDL-01 WT gave robust shutoff while WF10 WT gave poor shutoff which could be switched, significantly, upon the introduction of the reciprocal mutations at position 26 (Figure 5E, F).

Overall, these results show that UDL-01 has a PA-X capable of causing robust host shutoff in avian and mammalian cells while WF10 does not, and the reason for this difference maps to the identity of the amino acid at position 26.

### Differences in PA-X alone are not responsible for the attenuation of WF10 compared to UDL-01

The changes at PA position 26 are responsible for attenuation of WF10 *in vitro* and *in vivo*; these correlated better with the *in vitro* host cell shutoff activity of PA-X than with polymerase activity. To investigate whether the E26K polymorphism exerted its *in vivo* phenotypic effect via PA-X rather than through PA, we introduced a well characterized set of nucleotide substitutions into the frameshift site (FS mutant) of PA which has previously been shown to inhibit expression of PA-X [19, 34, 35]. We further combined this FS mutation with the reciprocal mutants at position 26.

We initially investigated the combined effect of position 26 and FS mutations on host shutoff in mammalian and avian cells (Figure 6A,B). For UDL-01 WT introduction of either the E26K or FS mutation individually or together ablated shutoff activity, indicating as we and others have shown, that segment 3 shutoff activity maps to PA-X [19, 34, 35, 37, 38]. Conversely, as WF10 WT segment 3 had no shutoff activity the FS mutant alone had no additional effect but ablated the increased shutoff activity seen when combined with WF10 K26E. An identical outcome of the mutations was observed in avian cells (Figure 6B). Finally, using whole viruses with both mutations at position 26 and FS combined we showed that when PA-X expression was ablated in both viruses, shutoff activity (as assayed by puromycin incorporation) was lost (Figure 6C). Overall, these data indicate the shutoff activity of segment 3s that contain PA 26E is entirely dependent on PA-X expression.

**Figure 6.**
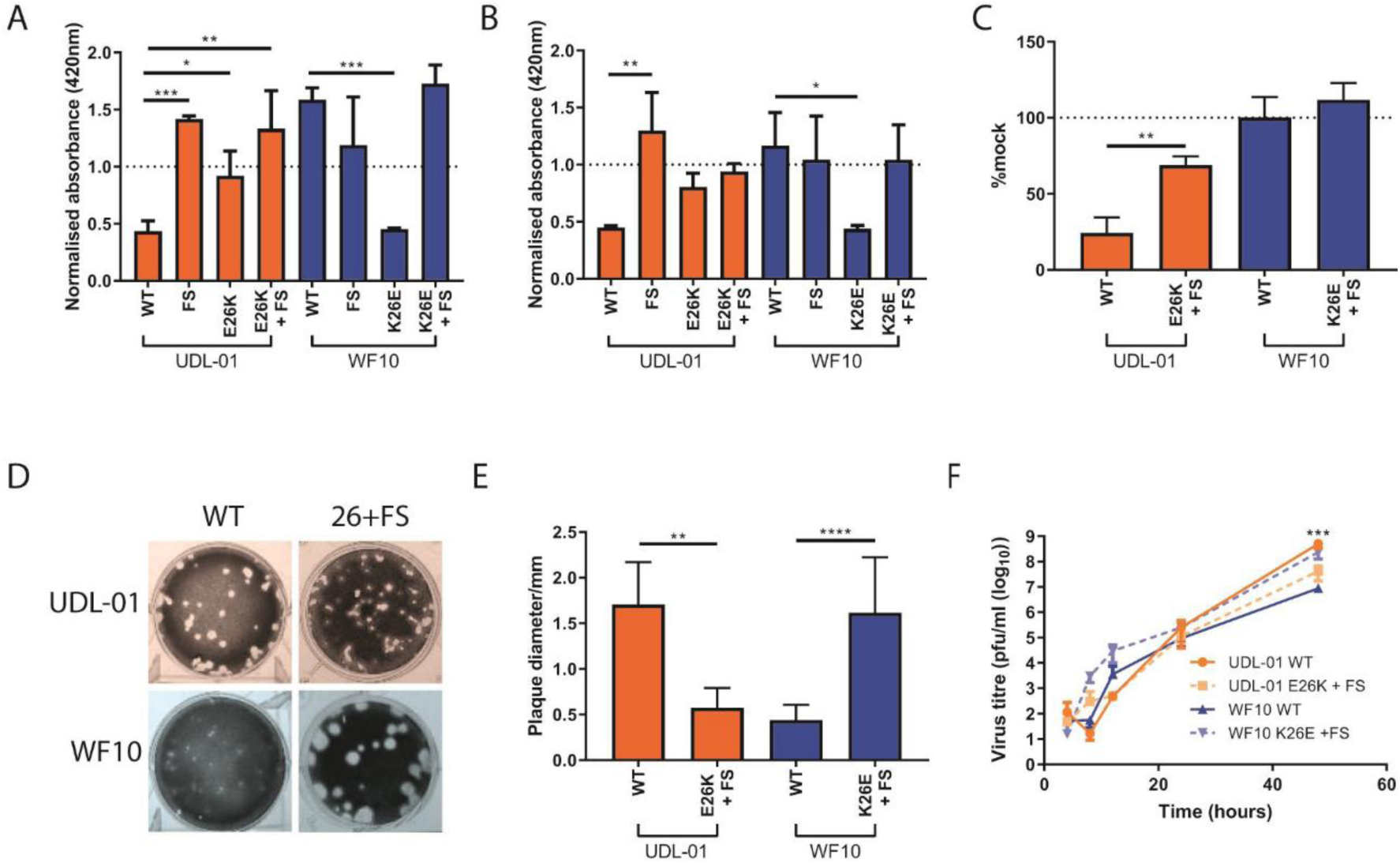
The attenuated replication of WF10 is independent of PA-X expression. H9N2 segment 3s with differing mutations were transfected into (A) 293T cells or (B) DF-1 cells along with a β-galactosidase (β-gal) reporter plasmid. 48 hours post transfection cells were lysed and then levels of β-gal assessed by colorimetric enzyme assay. Results were normalised to an empty vector control where no shut off host gene expression was expected. Graph represents the average of 3 independent experiments +/- SD., P values: ***= 0.0001; **= <0.0097 * = <0.045 (one-way ANOVA – 293T all and DF-1 WF10 panel or Kruskal Wallis – DF-1 UDL-01 panel with multiple comparisons). (C) Host cell shut off within viral infection was then assessed, MDCK cells were infected with each virus at a high MOI (5). 7 hours post-infection cells were pulsed with puromycin for 30 minutes. Cells were lysed in SDS-PAGE buffer and western blotted for puromycin. The area under the curve for each section was calculated and then compared to the mock infected sample. The Graph displays average +/- SD of 3 independent experiments. P values = **: 0.0042 (unpaired T-test). (D,E) The plaque phenotype of viruses containing both polymorphisms at position 26 and a PA-X frameshift mutation was assessed. Viral plaque phenotypes were assessed in MDCK cells under 0.6% agarose. 48 hours post-infection cells were fixed and stained with 0.1% crystal violet solution and (D) plaques imaged. (E) ImageJ analysis software was used to measure the diameters of 20 plaques per virus. Graph represents average +/- SD. P values = **:0.0015, ****=<0.0001 (unpaired T-test). (F) MDCK cells were infected with a low MOI of virus (0.01), cell supernatants were harvested at various time points post-infection and viral titres determined via plaque assay. Graph represents the average of 3 independent experiments +/- SD. P values =, ***= 0.006 (unpaired T-test – UDL-01 4 and 24 h.p.i. WF10 8, 12, 24, 48 h.p.i. or Mann Whitney – UDL-01 8, 12 and 48 h.p.i. WF10 4 h.p.i.). *** = between WF10 WT and WF10 K26E +FS.

We next tested whether the difference in plaque and replication phenotype also mapped to PA-X and whether the differences seen previously in this study were sensitive to removal of PA-X expression. Looking at the plaque phenotypes of these combined mutants we saw the frameshift mutant had little or no effect on the reciprocal plaque sizes seen in the viruses; although the UDL-01 E26K + FS mutant had a significantly smaller plaque size than UDL-01 WT, the WF10 K26E + FS virus retained its large plaque size, despite a lack of PA-X expression and shutoff activity (Figure 6D, E). Furthermore, the replication kinetics of the combined mutant viruses exhibited a similar phenotype – WF10 K26E + FS grew to higher titres than WF10 WT, indicating again that the enhanced replication conferred by the PA K26E mutation was independent of PA-X expression and shutoff activity (Figure 6F). Overall this implies that the difference in virus replication, and potentially pathogenicity, seen in viruses with differences at segment 3 position 26, may partially or fully map to PA, rather than PA-X alone.

## Discussion

In this study we investigated how differences in the PA gene of a progenitor (WF10) and a contemporary reassortant (UDL-01) H9N2 viruses led to differences in replicative fitness. We mapped these differences to a single PA residue at position 26, which is within the endonuclease domain. Although changes at this residue did not affect virus polymerase activity, they did cause reciprocal differences in replicative fitness in both mammalian and avian systems, in cell lines, primary cells, embryonated eggs, as well as *in vivo*, in chickens. We found that although these mutations strongly affected the shutoff activity of the accessory protein PA-X, this did not explain the differences in *in vitro* virus replication phenotype, indicating that it is likely that differences in PA function are partially, or fully, responsible for this.

The influenza virus accessory protein, PA-X, has been described in multiple studies as a virulence factor in avian influenza viruses (including H9N2 viruses) that can affect disease outcome in mammals or birds [22, 24, 26, 34]. Although other studies have found PA-X expression can lead to an attenuated phenotype, particularly in highly pathogenic H5N1 viruses [23, 25]. We found that differences between UDL-01 and WF10 at position 26 do modulate PA-X shutoff activity; however, PA-X activity alone was not responsible for the different replication phenotypes seen in these viruses, it is possible PA-X activity may still be contributing *in vivo* but that this effect is overshadowed by a dominant PA-specific replication effect. Although several further gene products are described as being generated from influenza A virus segment 3 (for example PA-N155 and PA-N182), these products do not share the PA endo domain and therefore are unlikely to explain the difference in phenotype between UDL-01 and WF10 [39].

The poor replication and small plaque phenotype of WF10 has been previously described by Wan and colleagues; the authors showed that the small plaque phenotype of WF10 could be overcome by supplying the virus with the internal genes of a human H3N2 virus [40]. In our study we further map these phenotypes to a single polymorphism in the PA gene at position 26. In a separate study by Obadan and colleagues, a WF10 mutant virus library with heterogeneity in the haemagglutinin receptor binding site was used to infect quails. It was found that PA-K26E, the UDL-01-like mutation, was consistently found to spontaneously arise – further suggesting that PA-K26 is responsible for the attenuated phenotype seen in WF10, both *in vitro* and *in vivo* [41].

Throughout this study it has been shown that PA residue 26 is responsible for the attenuated phenotype seen in WF10. When the relative distribution of polymorphisms at position 26 is looked at in the population of H9N2 viruses, or throughout avian influenza viruses in general, it becomes clear that the WF10-like K26 is very rare, with only a handful of viruses sharing any kind of polymorphism at this position. Over 99% of avian influenza viruses, including strains of the H5, H7 or H9 subtype contain the UDL-01-like E26 at this position, with a very few viruses containing lysine, glycine, aspartic acid or glutamine (Table 2).

**Table 2.** Prevalence of PA residue 26 polymorphisms in avian influenza viruses.

| Viruses | Glutamic acid (%) | Lysine (%) | Other (%) |
| --- | --- | --- | --- |
| All avian influenza viruses | 99.92 | 0.02 | 0.04 (glycine), 0.01 (aspartic acid), 0.01 (glutamine) |
| H5Nx | 99.73 | 0.03 | 0.07 (glycine), 0.11 (aspartic acid), 0.03 (glutamine) |
| H7Nx | 99.92 | 0.00 | 0.08 (glycine) |
| H9Nx | 99.87 | 0.13 | 0.00 |

Understanding the molecular basis of increased fitness of avian influenza viruses, both in avian and mammalian cells, as they continue to circulate and adapt to avian hosts is key to assessing the threat these viruses pose to food systems and to the human population. In this study we describe a single naturally occurring polymorphism in the endo domain of PA that leads to an attenuated replication and virulence phenotype. This work will help guide future surveillance efforts and may help us better understand the molecular basis of viral fitness and virulence in the avian host.

## Materials and Methods

### Ethics Statement

All animal experiments were carried out in strict accordance with the European and United Kingdom Home Office Regulations and the Animal (Scientific Procedures) Act 1986 Amendment regulation 2012, under the authority of a United Kingdom Home Office Licence (Project License Numbers: P68D44CF4 X and PPL3002952).

### Cell lines

Madin_Darby canine kidney (MDCK) cells, Human embryonic kidney (HEK) 293T cells and chicken DF-1 cells were maintained in Dulbecco’s Modified Eagle Medium (DMEM; Sigma) supplemented with 10% (v/v) FBS and 100U/ml Penicillin-Streptomycin (complete DMEM). All cells were grown at 37°C, 5% CO_2_.

Primary chicken kidney (CK) cells were generated as previously described [42]. Briefly, kidneys from three-week-old specific pathogen free (SPF) Rhode Island Red breed birds were shredded, washed in PBS, trypsinised, then filtered. Cells were resuspended in CK growth media (EMEM + 0.6% w/v BSA, 10% v/v tryptose phosphate broth, 300U/ml penicillin/streptomycin), plated and grown at 37°C, 5% CO2.

Primary chicken embryo fibroblasts (CEFs) were generated from 10-day-old chicken embryos. Embryos were homogenised and treated with trypsin/EDTA solution. Supernatants were passed through a metal mesh filter and centrifuged to pellet cells. Cells were resuspended in CEF media (M199, 4% (v/v) FBS and 100U/ml Pen/Strep), plated and grown at 37°C, 5% CO2.

### Viruses and reverse genetics

A pair of H9N2 viruses were used through this study, A/chicken/Pakistan/UDL-01/2008 (UDL-01) and A/Guinea Fowl/Hong Kong/WF10/1999 (WF10). Both virus reverse genetics systems were created using the bi-directional PHW2000 plasmids [43, 44]. Mutant PA segments were generated by site directed mutagenesis or Gibson assembly (NEB).

Reverse genetics viruses were generated as previously described [43]. Briefly, 250ng of each plasmid for either UDL-01 and WF-10 viruses were co-transfected into 6 well plates of 293Ts using lipofectamine 2000. 16h post transfection, media was changed to reverse genetics media (DMEM + 2mM glutamine, 100U/ml penicillin, 100U/ml streptomycin, 0.14% (w/v) BSA, 5µg/ml TPCK-treated trypsin). Following a 48h incubation at 37°C, 5%CO_2_ supernatants were collected and inoculated into embryonated hens’ eggs to grow virus stocks.

### Virus plaque assays

All plaque assays were performed in MDCK cells using 0.6% agarose overlay. Cells were stained with either 0.1% crystal violet solution (20% methanol). When plaques needed to be visualised by immunofluorescence, cells were fixed with 10% neutral buffered formalin followed by permeabilisation with PBS (0.2 %Triton X-100) then incubated with mouse monoclonal α-NP (Iqbal laboratory, 1:2000). Primary antibody was detected with goat anti-mouse IgG 568 conjugated secondary antibody (LICOR; 1:10000) Plates were imaged using an Odyssey Clx Near-Infrared Fluorescence Imaging System (LICOR). ImageJ was used to measure and analyse plaques.

### Minireplicon assays

DF-1 cells were seeded into 24 well plates were co-transfected with PB2, PB1, PA and NP, along with a firefly luciferase reporter construct under an avian polI promoter (CKpPol I Luc) at the following concentrations: PB2-160ng, PB1-160ng, PA-40ng, NP-320ng, pPol I Luc-160ng. 48 hours post-transfection cells were lysed in Passive Lysis Buffer (Promega) and lysates were read on a Promega GloMax Multi Detection unit using Luciferase Assay Reagent II (Promega) following the manufacturer’s instructions.

### Virus replication *in vitro* and *in ovo*

MDCK and CK cells were inoculated with virus diluted in serum free DMEM for 1 h at 37°C at an MOI of 0.01. Cell supernatants were taken at 4-, 8-, 12-, 24-, 48- and 72-hours post-infection. After 1 hour incubation with virus, cells were then washed twice to remove unbound virus, and media was replaced with virus growth medium - DMEM plus 2 μg/ml tosyl phenylalanyl chloromethyl ketone (TPCK)-treated trypsin for MDCK cells or Eagle’s minimum essential medium (EMEM), 7% bovine serum albumin and 10% tryptose phosphate broth for CKCs.. Viruses were titred by plaque assay on MDCK cells.

10-day-old embryonated hens’ eggs (VALO breed) were inoculated with 100 pfu of virus into the allantoic cavity. Eggs were incubated for 4-72h and culled via the schedule one method of refrigeration at 4°C for a minimum of 6h. Harvested allantoic fluid from each egg was collected and clarified by centrifugation, virus titres were assessed by plaque assay on MDCKs.

### Virus infection, transmission and clinical outcome *in vivo*

Prior to the commencement of the study, all birds were swabbed (in both oropharyngeal and cloacal cavities) and bled via wing prick to confirm they were naïve to the virus. All infection experiments were performed in self-contained BioFlex B50 Rigid Body Poultry isolators (Bell Isolation Systems) at negative pressure. 20 birds per group were directly inoculated with 10^4^ pfu of virus intranasally. Mock infected birds were instead inoculated with sterile PBS. One day post-inoculation 8 naïve contact birds were introduced into each isolator to determine virus transmission.

Throughout the experiment, birds were swabbed in the buccal and cloacal cavities (days 1-8, 10 and 14 post-infection). Swabs were collected into 1ml of virus transport media (WHO standard). Swabs were soaked in media and vortexed for 10 seconds before centrifugation. Viral titres in swabs were determined by plaque assay on MDCKs.

At day 1 and 3 post-inoculation, directly infected birds were euthanisedand a panel of tissues were collected and stored in RNA later at -80°C until further processing. On day 14 post-infection, all remaining birds were culled via overdose of pentobarbital or cervical dislocation.

### RNA extraction and RT-PCR from chicken tissues

30mg of tissue collected in RNA later was mixed with 750µl of Trizol. One sterile 5mm stainless steel bead was added per tube and tissues were homogenised using the Retsch MM 300 Bead Mill system (20Hz, 4 min). 200µl of chloroform was added per tube and tubes were shaken vigorously and incubated for 5 min at room temperature. Samples were centrifuged (9,200xg, 30 min, 4°C) and the top aqueous phase containing total RNA was added to a new microcentrifuge tube and the remaining fluid discarded. RNA extraction was then carried out using the QIAGEN RNeasy mini kit following manufacturers’ instructions.

100ng of RNA extracted from tissue samples was used for qRT-PCR. All qRT-PCR was completed using the Superscript III platinum One step qRT-PCR kit (Life Technologies) following manufacturer’s instructions for reaction set up. Cycling conditions were as follows: i) 5 min hold step at 50°C, ii) a 2 min hold step at 95°C, and 40 cycles of iii) 3 sec at 95 °C and iv) 30 sec annealing and extension at 60 °C. Cycle threshold (CT) values were obtained using 7500 softwarev2.3. Mean CT values were calculated from triplicate data. Within viral M segment qRT-PCR an M segment RNA standard curve was completed alongside the samples to quantify the amount of M gene RNA within the sample from the CT value. T7 RNA polymerase-derived transcripts from UDL-01 segment 7 were used for the preparation of the standard curve.

### Host shutoff assays

β-galactosidase (β-gal) shutoff reporter assays were performed as previously described [19]. Briefly, 293T or DF-1 cells were co-transfected with expression plasmids for the influenza segment 3 and β-gal reporter. 48 hours later, cells were lysed with Reporter lysis buffer (Promega). β-gal expression was measured using the β-galactosidase enzyme assay system (Promega). A Promega GloMax Multi Detection unit was employed to read absorbance at 420nm.

For the radio-labelling shutoff activity assays using live virus, chicken embryonic fibroblast (CEFs) were infected with 7:1 (PR8: H9N2) reassortant viruses containing the described PAs at an MOI of 3. At 6 hours post-infection, cells were washed and overlaid with 1ml of methionine-and cysteine-free DMEM supplemented with 5% dialysed FCS and 2mM L-Glutamine to starve the cells of methionine and cysteine. At 8 hours post-infection, cells were washed and overlaid with methionine- and cysteine-free DMEM (supplemented as above) including ^35^S-methionine/cysteine protein labelling mix (Perkin/Elmer) at 0.8mBq/ml. Cells were incubated at 37°C in a vented box containing activated charcoal (Fisher) for 1 hour. Cells were washed once with ice-cold PBS and then cells lysed in protein loading buffer for SDS-PAGE and processed via autoradiography. Gels were fixed in gel fix solution (50% methanol, 10% acetic acid) for 5-15 minutes. Fix solution was replaced for 2 more rounds of fixing. Gels were dried in a gel dryer (Bio-Rad) by heating up to 80°C for 2-4h under vacuum pressure. Dried gels were placed in a sealed cassette with an X-ray film (Thermo Fisher) overnight, at a minimum or until the desired signal strength was achieved. X-ray films were developed using a Konica SRX-101A X-ograph film processor using manufacturers’ instructions.

For the non-radioactive shutoff activity assays using live virus, MDCKs were infected with whole H9N2 virus at an MOI of 5. At 7.5 hours post-infection cells were washed and the medium changed to complete DMEM containing 10µg/ml of Puromycin dihydrochloride from *Streptomyces alboniger* for 30 minutes. Cells were washed then lysed in protein loading buffer for SDS-PAGE and western blotted, probing for puromycin. Puromycylated protein synthesis was quantified in the region of the gel between 45kDa and 80kDa.

Protein quantification following autoradiography or anti-puromycin western blot was determined by densitometry using ImageJ analysis software.

### Bioinformatics analysis

To assess the prevalence of different polymorphisms at position 26 of PA, every amino acid sequence of full length PA isolates from avian hosts, excluding duplicate sequences, was downloaded from the NCBI Influenza Virus Database (https://www.ncbi.nlm.nih.gov/genomes/FLU/Database/nph-select.cgi), as of the 23^rd^ May, 2020. Sequences were aligned using Geneious R11.1.5 and the distribution of different amino acids was recorded.

### Statistical analysis

All statistical analysis was carried out using GraphPad Prism 6/7 software. Distribution of data was assessed prior to deciding on the statistical test to use.

## Acknowledgments

We would like to thank the animal housing staff for looking after the wellbeing of chickens used in this study and for monitoring their health throughout the experiments.

